# DISMIR: a deep learning-based cancer-detection method by integrating DNA sequence and methylation information of individual cell-free DNA reads

**DOI:** 10.1101/2021.01.12.426440

**Authors:** Jiaqi Li, Lei Wei, Xianglin Zhang, Wei Zhang, Haochen Wang, Bixi Zhong, Zhen Xie, Hairong Lv, Xiaowo Wang

## Abstract

Detecting cancer signals in cell-free DNA (cfDNA) high-throughput sequencing data is emerging as a novel non-invasive cancer detection method. Due to the high cost of sequencing, it is crucial to make robust and precise prediction with low-depth cfDNA sequencing data. Here we propose a novel approach named DISMIR, which can provide ultrasensitive and robust cancer detection by integrating DNA sequence and methylation information in plasma cfDNA whole genome bisulfite sequencing (WGBS) data. DISMIR introduces a new feature termed as “*switching region*” to define cancer-specific differentially methylated regions, which can enrich the cancer-related signal at read-resolution. DISMIR applies a deep learning model to predict the source of every single read based on its DNA sequence and methylation state, and then predicts the risk that the plasma donor is suffering from cancer. DISMIR exhibited high accuracy and robustness on hepatocellular carcinoma detection by plasma cfDNA WGBS data even at ultra-low sequencing depths. Analysis showed that DISMIR tends to be insensitive to alterations of single CpG sites’ methylation states, which suggests DISMIR could resist to technical noise of WGBS. All these results showed DISMIR with the potential to be a precise and robust method for low-cost early cancer detection.

## INTRODUCTION

Cell-free DNA (cfDNA) are degraded DNA fragments released to body fluids such as plasma and urine mainly brought by apoptosis or necrosis cells (Crowley *et al.*, 2013). It was reported that in the early stage of cancer when there are no significant clinical symptoms on patients, the state of DNA in cancer cells has already changed (Baylin *et al.*, 2001) and can be detected in the plasma of cancer patients as circulating tumor DNA (ctDNA) (Schwarzenbach *et al.*, 2011). With the development of high-throughput sequencing technologies, non-invasive approaches by identifying cancer signals in cfDNA sequencing data are emerging as novel liquid biopsy methods for cancer diagnosis (Wan *et al.*, 2017).

The majority of cfDNA studies focus on the mutation of oncogenes. The existence and fraction of ctDNA in the total cfDNA is calculated by detecting certain mutations in a small oncogene panel (Bettegowda *et al.*, 2014; Abbosh *et al.*, 2017). However, the fraction of ctDNA in early-stage cancer is too low to detect without an ultra-deep sequencing method (Newman *et al.*, 2014; Heitzer *et al.*, 2018). Besides, mutations that drive carcinogenesis are usually diverse, leading to heterogeneity across different patients or across different loci in tumor tissues, which limits the potent of detecting cancer by ctDNA mutation (Burrell *et al.*, 2013). Some other studies tried to detect the rearrangement of chromosomes during carcinogenesis by cfDNA such as copy number alterations (Chicard *et al.*, 2016; Weiss *et al.*, 2017) and fragmentation patterns (Snyder *et al.*, 2016; Cristiano *et al.*, 2019), and found interesting relationships between these signatures and cancer. However, as cfDNA sequencing data are mixed data with low signal-to-noise ratios, these low-resolution signatures can hardly be distinguished from noise when detecting early-stage cancer, therefore cannot be solely applied as accurate biomarkers for early-stage cancer detection.

The methylation states of DNA are altered in the early stage of cancer widespread across the whole genome (Feinberg *et al.*, 2006; Alvarez *et al.*, 2011), which warranties methylation as an informative feature for early-stage cancer detection. Therefore, integrations of the methylation states on different CpG sites (Warton and Samimi, 2015) or in different sub-genomic regions (Chan *et al.*, 2013) are promising approaches to enhance the precision of cancer detection. Furthermore, as the fraction of ctDNA in the total cfDNA was shown to be concordant with tumor burden (Adalsteinsson *et al.*, 2017), deconvolution of cfDNA to infer its origin becomes a hopeful approach to estimate the existence and severity of cancer. For example, Feng *et al*. used the methylation ratio of certain regions to do deconvolution with non-negative matrix factorization or quadratic programming (Feng *et al.*, 2019). Though, the performance of such methods is still limited by the low signal-to-noise ratio.

Recently, an approach called CancerDetector proposed to predict the source of cfDNA at the resolution of individual sequencing reads using the local correlation of methylation states between adjacent CpG sites (Li *et al.*, 2018), providing a novel read-based sight to investigate cfDNA sequencing data. However, different depths of sequencing data may introduce systematic deviation to the prediction results of CancerDetector, which could further reduce the accuracy of cancer diagnosis.

Previous work suggested that the methylation states are partly *cis*-regulated by the surrounding DNA sequence (Lienert *et al.*, 2011; Cedar and Bergman, 2012). Therefore, the surrounding DNA sequence may provide valuable information to analyze the methylation state and predict the source of individual reads. Here, we adopted a deep learning model named DISMIR to predict the source of individual reads. DISMIR can integrate the DNA sequence and methylation information of the selected differentially methylated regions (DMRs) across the whole genome, and thus enables the prediction accuracy even at very low sequencing depths. Besides, we introduced a new feature termed as “*switching region*” to find specific DMRs suitable for source prediction of individual reads to further improve the accuracy. DISMIR successfully achieved an area under the receiver operating characteristic curve (AUC) of 0.9932 ± 0.0038 (mean ± SD) in the diagnosis of hepatocellular carcinoma (HCC) with low sequencing depth cfDNA whole-genome bisulfite sequencing (WGBS) data (coverage from 1× to 3×). When subsampling the sequencing data to an ultra-low sequencing depth (from 0.01× to 0.03×), DISMIR still achieved an AUC of 0.9033 ± 0.0396. Analysis of the deep learning model showed that DISMIR successfully extracted DNA sequence and methylation patterns related to HCC across the whole genome, and was more sensitive to global methylation alterations, which made DISMIR able to resist to technical noise of WGBS. The results suggested DISMIR can do better cancer diagnosis with low sequencing depths at the early stage of cancer by successfully combining the information of DNA sequence and methylation together, which could be of great help to further clinical application.

## MATERIAL AND METHODS

### Overview

The ultimate goal of DISMIR is to diagnose cancer by integrating DNA sequence and methylation information in plasma cfDNA WGBS data. The diagnosis is performed by predicting the source of each read and then estimating the proportion of tumor-derived reads in the total cfDNA. The overall procedure of DISMIR comprises four main steps: (1) Identify the cancer-specific differentially methylated regions (DMRs) of cancer tissues in comparison with healthy people’s plasma across the whole genome as candidate biomarkers (Figure 1A). (2) Screen out reads in plasma cfDNA WGBS data that are located in the cancer-specific DMRs (Figure 1B). (3) Train a deep learning model to integrate DNA sequence and methylation information with these data to mark each read a value named d-score as the potent that the read is derived from cancer tissues (Figure 1C). (4) Estimate the fraction of tumor-derived reads of a plasma sample by all d-scores to infer whether the plasma donor is suffering from cancer (Figure 1D). Here we adopted hepatocellular carcinoma (HCC) as an example to validate the performance of DISMIR.

**Figure 1.**
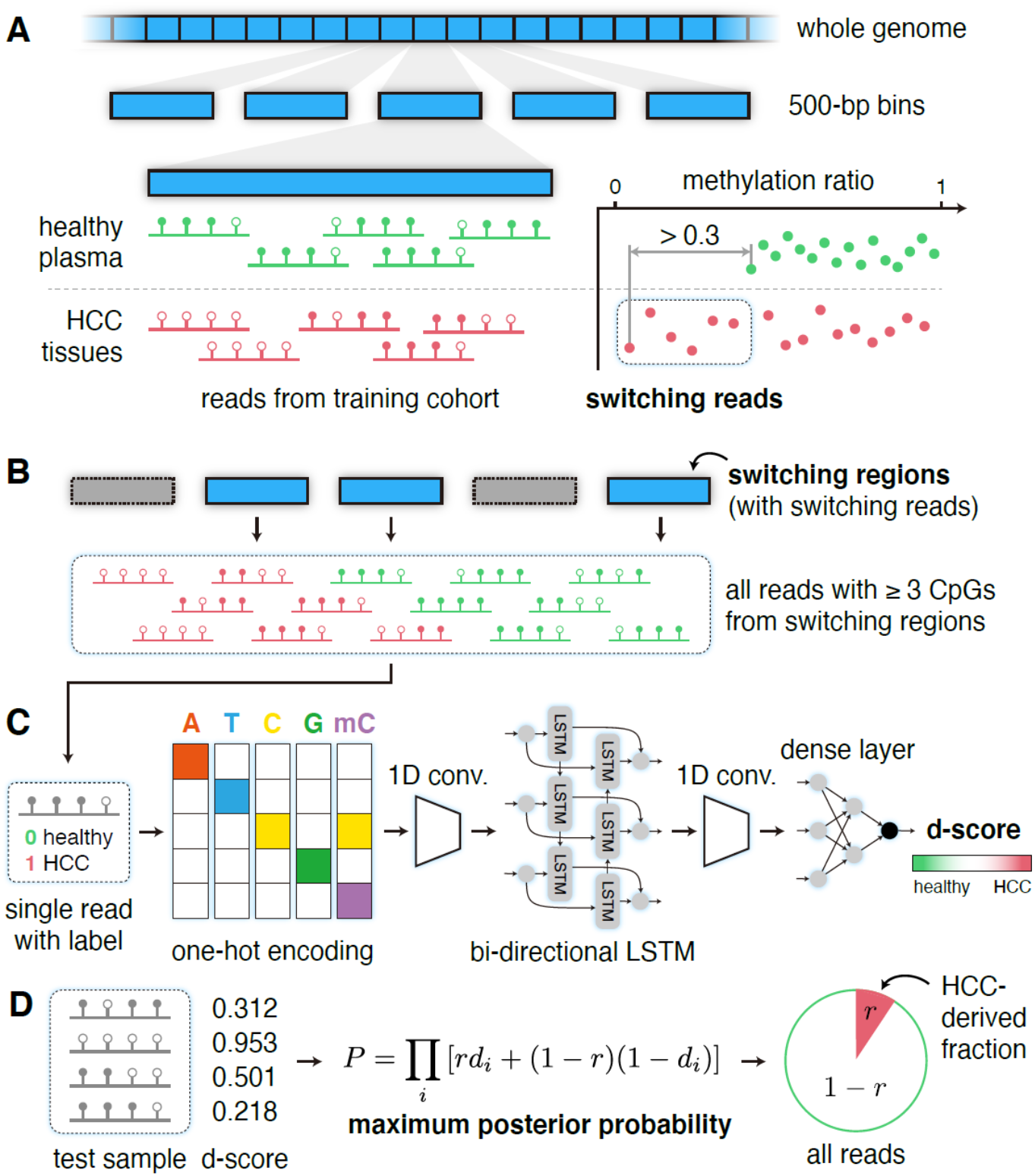
Overview of DISMIR. (A) Identifying cancer-specific DMRs across the whole genome with the definition of switching regions and switching reads. (B) Collecting all reads with 3 or more CpG sites from switching regions for further analysis. (C) Training a deep learning model to calculate the d-score of each individual read. (D) Predicting tumor fraction of a sample by maximizing the posterior probability.

### Data collection and processing

The data employed in this study contains single-end WGBS data of plasma cfDNA from 32 healthy people, 8 hepatitis B virus (HBV) carriers without cancer and 24 HCC patients with European Genome-Phenome Archive database (EGA) accession number EGAS00001000566 (Chan *et al.*, 2013). Among them, 13 HCC patients have single-end WGBS data of cancer tissues paired with plasma cfDNA. The training cohort contains 9 HCC patients’ cancer tissues WGBS data and 18 randomly chosen healthy people’s plasma WGBS data (randomly chosen for 10 times). WGBS data of the remaining 14 healthy people’s plasma, 8 HBV carriers’ plasma and 11 unpaired HCC patients’ plasma compose the test cohort. The WGBS data of the remained 4 HCC tissues were used for simulation experiments to evaluate the effect of our approach.

We used BS-Seeker2 (Guo *et al.*, 2013) to align all these WGBS data to hg19, removed PCR duplicates and then called the methylation states of all CpG sites for subsequent analysis.

### Identifying HCC-specific differentially methylated regions

Identifying HCC-specific DMRs across the whole genome could provide valid cancer-related information refraining from the unconcerned variation of methylation states among different samples. As the sequencing data of plasma cfDNA could be regarded as a mixed signal of tumor-derived cfDNA and basal cfDNA, which is similar to cfDNA at the healthy state, we should use the reads from regions where the methylation patterns are different between cancer tissues and healthy plasma cfDNA. Previous studies (Liggett *et al.*, 2010; Jühling *et al.*, 2016; Li *et al.*, 2013; Hebestreit *et al.*, 2013; Wu *et al.*, 2015) have produced many methods to define DMRs. These methods mainly focused on the statistics of total reads from a certain genome region. However, fractions of tumor-derived reads in plasma cfDNA are usually ultra-low especially at the early stage. The identification of tumor-derived reads would be greatly dampened by outliers from healthy tissues in the calculation of traditional statistics. Therefore, we defined DMRs as regions where the methylation patterns of tumor-derived reads are distinguishable from patterns of reads from healthy plasma to enhance the cancer-related signal at read-resolution in cfDNA sequencing data.

Based on such assumption, we introduced a new feature named “*switching regions*” and “*switching reads*”, which were defined with the following steps (Figure 1A). Firstly, we divided the whole genome into 500-bp regions without overlaps and filtered out regions with less than 25 reads in all training-cohort samples. Then we calculated the methylation ratios of all DNA fragments from a certain region to get their distributions in cancer tissues as well as cfDNA from healthy plasma. Here we only used reads with three or more CpG sites. Next, we compared the maximum and minimum values of two distributions. For instance, HCC shows a significant genome-wide hypomethylation pattern in comparison with healthy tissues (Chan *et al.*, 2013), so we focused on the hypomethylated regions in HCC. Here we denoted the healthy plasma’s minimum methylation rate in a region as *H*_min_, and denoted the cancer tissues’ minimum methylation rate as *T*_min_. When *H*_min_ – *T*_min_ is larger than a certain threshold (set to 0.3 in this study), this region is defined as a switching region. All reads from switching regions with methylation rates lower than *H*_min_ are defined as switching reads.

### Predicting the source of each read with a deep learning model

To gain a valid and comprehensive model to depict the DNA sequence and methylation pattern in tumor-derived reads in cfDNA WGBS data, we built a deep learning model to predict the potent that a read is derived from cancer tissues, termed as d-score. All reads with three or more CpG sites from switching regions were used to train the deep learning model. By attaching label to each read according to its source (from healthy plasma as 0, from cancer tissues as 1), we converted this problem into a binary classification problem of reads. For each read, the first 5 bp at the 5’ end was trimmed to avoid the influence of adapters. Then all reads were trimmed at the 3’ end to a same length (*L* = 66 in this study) to unify the input format. We randomly subsampled the reads to ensure the balance between the amount of two sample types and reserved 20% of these reads for kernel visualization.

Here we referred to the structure of DanQ model (Quang and Xie, 2016), which was built to quantify the function of DNA sequences, and made some adjustments on it to serve as the core of the deep learning model (Figure 1C). Each base of a unified read was encoded into a one-hot matrix according to the nucleobase, and the methylation state of the base was also encoded, where 1 presents methylated and 0 presents unmethylated. Therefore, each input read was encoded into a *L*×5 matrix. After the input layer, we sequentially added a one-dimensional (1D) convolution layer, a maxpooling layer, a bi-directional LSTM layer, a 1D convolution layer, a flatten layer, and three dense layers. The output of the model was a continuous value denoted as d-score between 0 and 1 corresponding to the label of each read. The closer the d-score is to 1, the more likely the read is from a cancer tissue.

### Estimating the fraction of tumor-derived cfDNA

The d-score calculated by the deep learning model was treated as the probability that the read is from a cancer tissue. For a tested sample with *n* reads and their d-scores *d*_1_, *d*_2_, … *d*_*n*_ (Figure 1D), we inferred the proportion of reads from cancer tissue according to these d-scores by calculating the maximum posterior probability inspired by CancerDetector (Li *et al.*, 2018). When given the ratio of tumor-derived reads as *r* for a sample, and assuming that d-scores of each read are independent, we could get the posterior probability of this sample with these d-scores as *P*:

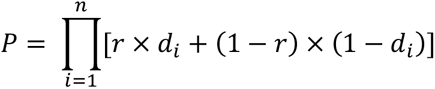

When maximizing *P*, we could get the estimated ratio of tumor-derived reads by DISMIR denoted as 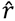, which could be regarded as the risk that the plasma donor is suffering from cancer for cancer diagnosis:

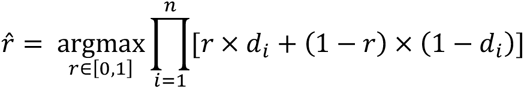

### Visualizing kernels of the deep learning model by position frequency matrices (PFM)

We visualized kernels of the deep learning model to figure out the DNA sequence and methylation patterns that the model focused on. After finishing the training of the deep learning model, we took out the weight matrices of all kernels in the first 1D convolution layer. We then used the reserved 20% reads in the training set on every possible position as inputs and calculated their activation values by the weight matrices. For each weight matrix, we located the top 1% output values and got the corresponding inputs. These inputs were superposed together with their activation values as weights to calculate the frequency of each base and the methylation state as PFMs.

## RESULTS

### DISMIR achieved high precision in early-stage HCC detection

We identified switching regions and trained DISMIR on a randomly selected training cohort, and then tested DISMIR on remained samples as the test cohort for ten times (see Material and Methods for details). For every random selection, we trained DISMIR for ten times with the same data and then applied the model on the test cohort. The average d-score of each individual read was calculated as the final d-score for downstream estimation of tumor fraction to get rid of the randomness of the deep learning method.

We adopted the receiver operating characteristic (ROC) curve to evaluate the ability of the tumor fraction predicted by DISMIR for distinguishing HCC patients from healthy people. As shown in Figure 2A, the area under the ROC curve (AUC) of DISMIR was 0.9932 ± 0.0038 (mean ± SD). At the specificity of 100%, DISMIR achieved a sensitivity of 90.91% ± 0.00%; while at the sensitivity of 100%, the specificity of DISMIR was 91.82% ± 4.69%. We further compared our approach with CancerDetector (Li *et al.*, 2018). As reported in the result of CancerDetector, the AUC during the detection of HCC with ten runs was 0.990 ± 0.004. At the specificity of 100%, CancerDetector had a sensitivity of 94.9% ± 2.7%. At the sensitivity of 100%, the specificity of CancerDetector was 77.3% ± 9.4%. To evaluate a more unbiased performance of CancerDetector, we further applied CancerDetector on the test cohort of this study (Figure 2A), and found that CancerDetector achieved an AUC of 0.9678 ± 0.0170. At the specificity of 100%, the sensitivity was 58.18% ± 23.16%. At the sensitivity of 100%, the specificity was 83.64% ± 4.39%. The results suggested that the effect of DISMIR was comparable with CancerDetector.

**Figure 2.**
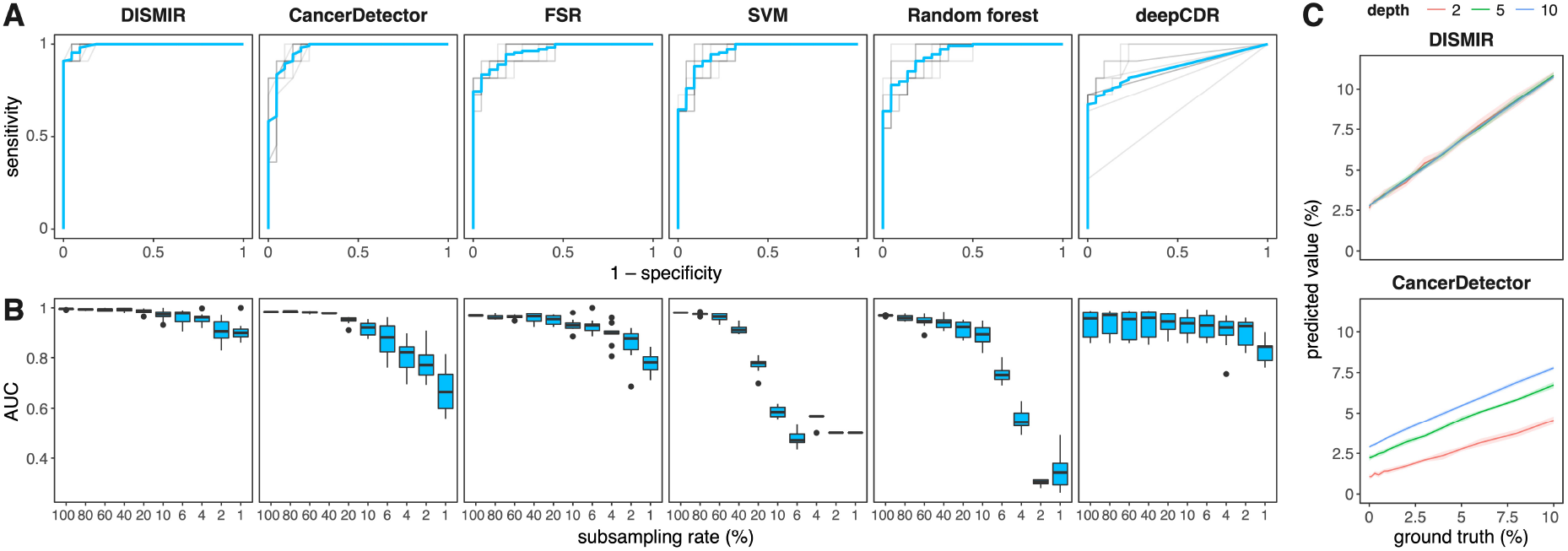
Results of DISMIR and other methods on HCC diagnosis. (A) ROC curves of different HCC-diagnosis methods in the test cohort. Blue lines show the average of the ROC curve. Each method was performed for ten times with random partition of training and test samples. (B) AUCs of different HCC-diagnosis methods at different subsampling rates. Each condition was performed for ten times with randomly subsampling in the test cohort. (C) Simulation results at different depths with DISMIR (top) and CancerDetector (bottom). Each condition was performed for ten times with randomly sampling and mixing.

To assess the contribution of the deep learning model, we adopted the fraction of switching reads (FSR in short) and trained two traditional machine learning models, SVM and random forests, based on the methylation ratios of each switching region to diagnose HCC. As shown in Figure 2A, these methods showed moderate classification accuracies but could hardly serve as effective HCC-diagnosis markers in comparison with DISMIR. Besides, we used reads from DMRs identified by CancerDetector, which had a similar coverage on genome with DMRs defined by switching regions, to train the same deep learning model (deepCDR in short), and found a lower precision than DISMIR (Figure 2A), further advocating the advantage of defining DMRs by switching regions. As a result, both the deep learning model and the definition of switching reads contributed to the great performance of DISMIR.

### Subsampling and simulation results showed DISMIR as an ultrasensitive and robust HCC detection method

To evaluate the performance of DISMIR at low sequencing depths, we randomly subsampled data in the test cohort for ten times, and applied DISMIR and other above-mentioned methods on these data. As shown in Figure 2B, DISMIR kept high precisions, while the accuracy of CancerDetector decreased significantly with the reduction of sequencing depths. When the data were subsampled with a ratio of 1% (coverage from 0.01× to 0.03×), DISMIR still achieved an AUC of 0.9033 ± 0.0396, which is significantly higher than the AUC of CancerDetector (0.6696 ± 0.0874). Interestingly, FSR exhibited higher AUCs at low sequencing depths than CancerDetector, suggesting that defining DMRs by switching regions could resist to noise better than traditional methods. What’s more, deepCDR also showed better performance at low sequencing depths than CancerDetector, which demonstrated the benefit of employing the deep learning model. Meanwhile, accuracies of the two traditional machine learning methods decreased rapidly with the sequencing depth reduction and even lost the discrimination ability when the subsampling ratio was less than 4% (Figure 2B). All the results suggested that learning the joint patterns of DNA sequence and methylation of reads from switching regions by the deep learning model could predict the source of reads more precisely and thus guarantees the sensitivity of HCC diagnosis at ultra-low sequencing depths.

We further conducted a simulated dataset to validate the robustness of DISMIR. We randomly sampled reads from WGBS data of HCC tissues and healthy plasma cfDNA respectively and mixed them together with different proportions to imitate certain tumor fractions. Besides, the total amount of reads also varied to simulate different sequencing depths. The sampling procedure was repeated for ten times for each condition. We then tested DISMIR and CancerDetector on the simulated dataset. As shown in Figure 2C, the predicted tumor fractions of DISMIR were consistent at different sequencing depths, but those of CancerDetector increased significantly with the increase of sequencing depth, which may introduce bias into the HCC-diagnosis approach as the sequencing depths can hardly be exactly the same without a loss-of-information subsampling procedure. The results suggested that DISMIR is highly robust at different sequencing depths and thus is more applicable.

### Kernels of DISMIR paid attention to joint patterns of DNA sequence and methylation

To investigate how DISMIR distinguished HCC-derived cfDNA fragments from others by employing DNA sequence and methylation information, we tried to interpret the deep learning model of DISMIR by investigating the network details. We visualized the kernels of the first 1D convolution layer by calculating their position frequency matrices (PFMs, see Methods and Materials for details). We compared the sequence patterns of these PFMs with known motifs by TOMTOM (Gupta *et al.*, 2007) and merged the *E*-values assigned by TOMTOM from ten times of training with Fisher’s combined probability test. 28 motifs were identified as significant motifs (*p*-value < 0.05) matching with the kernel PFMs (Supplemental Table 1). Interestingly, many of these motifs were related to HCC (Supplemental Table 1). For example, as shown in Figure 3A, two kernels were matched to the EGR2 and ZFP64 (ZF64A) motif respectively. EGR2 is an anti-tumor transcriptional factor, the induction of which could suppress the malignancy of HCC (Zeng *et al.*, 2017; Wang *et al.*, 2020). Meanwhile, the expression of ZFP64 was shown to be positively correlated to the overall survival of advanced HCC patients with the treatment of a second-line therapy (Bitzer *et al.*, 2016).

**Figure 3.**
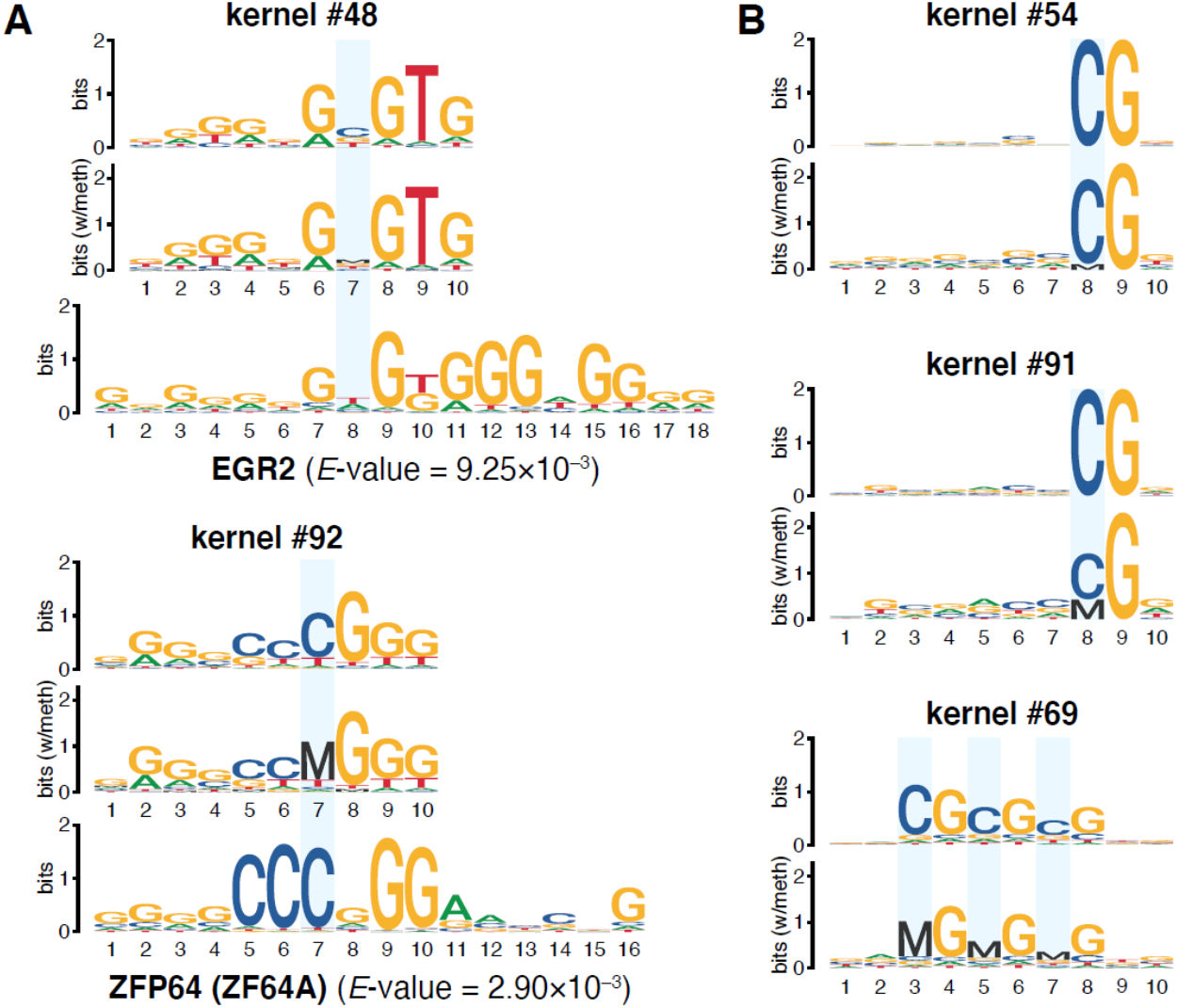
Visualized convolution kernels of the deep learning model. (A) Visualized kernels matched to known HCC-related motifs. (B) Visualized kernels focusing on the methylation state of CpG sites. Cytosines at CpG sites are marked with blue rectangles.

We further visualized the kernels with the methylation information. We treated the methylated cytosine (noted as ‘M’) and the unmethylated cytosine (noted as ‘C’) as two different base and then visualized the kernels in five-base logos. The results were similar with four-base logos except for significant difference at CpG sites. By such visualization, we successfully found the evidence that the deep learning model combined sequence with methylation information together. As shown in Figure 3A, the cytosine at the CpG site of the ZFP64-like kernel was almost fully methylated, suggesting the methylation state on this motif was highly coordinated with its flanking DNA sequence pattern during HCC detection. What’s more, we also found several kernels concentrated to different methylation states of CpG sites at different positions of reads (Figure 3B). For example, both kernel #54 and #91 paid attention to the CpG site at the 8^th^ position, but they attached quite different importance to the methylation state of the CpG site (Figure 3B). Therefore, with other kernels that might pay more attention to the information of joint patterns of DNA sequence and interior methylation, the deep network could thus combine the information together as the preliminary pattern extraction of a whole read for further analysis to predict the source of the read more accurately.

### DISMIR employed the joint pattern of DNA sequence and methylation to distinguish HCC-derived reads

Though kernel visualization suggested both DNA sequence and methylation information were processed in DISMIR, less was known about whether DNA sequence and methylation decided the results jointly. Therefore, we investigated the relationship between the methylation ratios of all reads and their d-scores derived by DISMIR (Figure 4A), which showed a significant negative correlation (Pearson’s *r* = –0.900). However, d-scores of reads with similar methylation ratios varied enormously. If the score of each read was assigned by its methylation ratio, the correlation should be much higher. Besides, as the methylation states of CpG dyads on both strands are correlated but could be different in some conditions (Shao *et al.*, 2009), we generated reverse complementary (RC) reads of raw reads from cancer tissues that were not in the training set with the same methylation state at every CpG dyad, thus the paired raw read and RC read shared the same methylation ratio. We then used DISMIR to predict the d-scores of raw reads and RC reads (Figure 4B). The results suggested that DISMIR successfully found the correlated pattern of reads derived from different strands (Pearson’s *r* = 0.869), while there exhibited some difference between them. All the results showed that DISMIR determined the d-score of a read by more beyond its methylation ratio.

**Figure 4.**
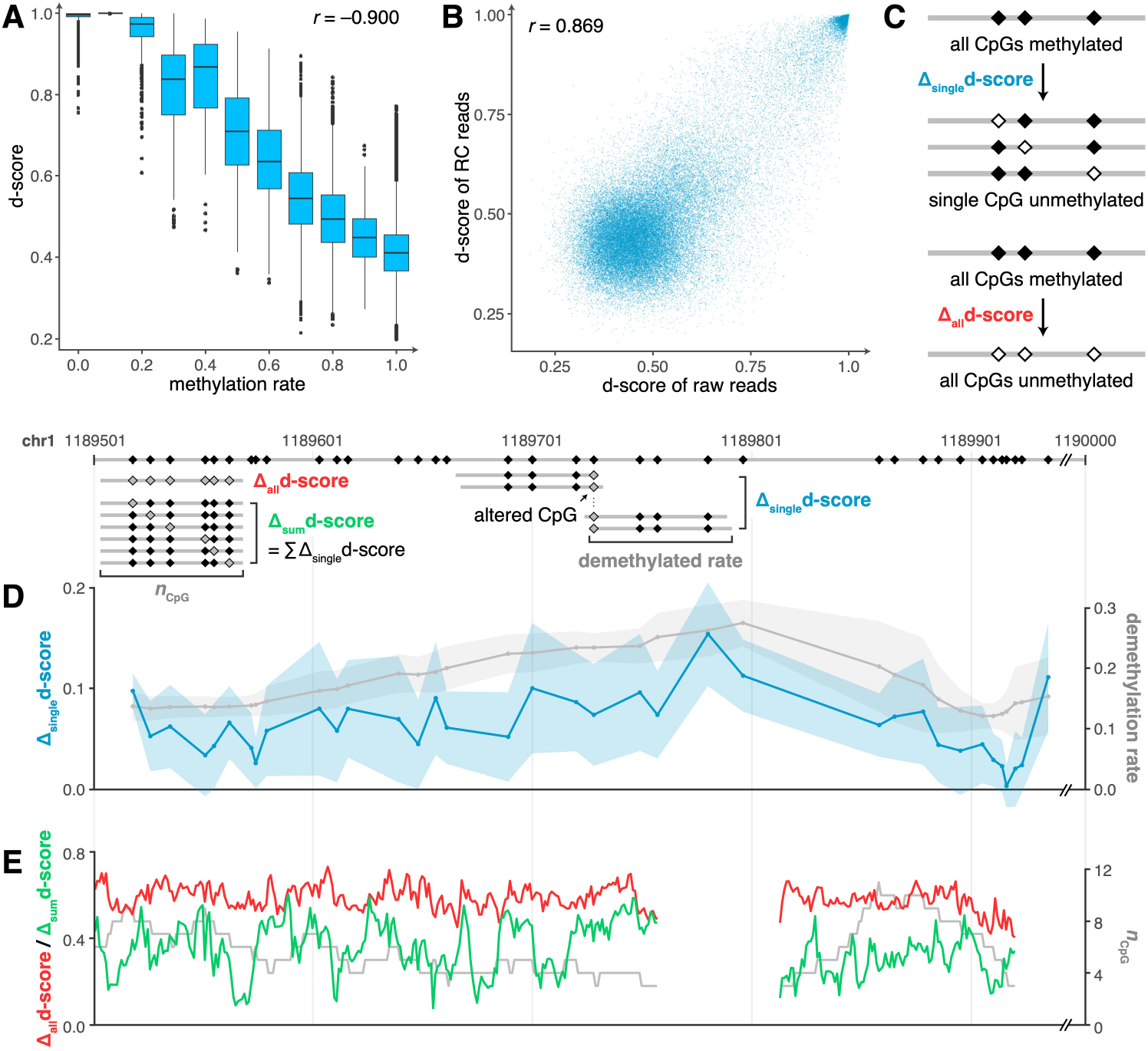
The joint pattern of DNA sequence and methylation decides the prediction of DISMIR. (A) The relationship between methylation rates and d-scores of all reads. (B) The relationship of d-scores of raw reads and their reverse complementary (RC) reads. (C) The schematic diagram depicting how reads with altered methylation states were generated for downstream analysis. (D) Δ_single_d-score (blue line) and the corresponding demethylation rate (gray line) at different positions in the selected switching region. Colored shadows show the standard deviation of Δ_single_d-score. (E) Δ_all_d-score (red line), Δ_sum_d-score (green line) and the corresponding CpG count (gray line) at different positions in the selected switching region.

Thus, we investigated the collaboration of DNA sequence and methylation within switching regions. Here we gave a sample from a certain switching region with a length of 500 bp on chromosome 1. We generated all possible reads with the same length as the input reads with three or more CpG sites within the region. All CpG sites on these reads were set to be methylated, and their d-scores were calculated by DISMIR. Then, the methylation state of each single CpG site on all reads was altered to be unmethylated. The d-scores changed correspondingly with a magnitude denoted as Δ_single_d-score (Figure 4C). Similarly, we examined reads with all CpG sites altered to the unmethylated state, and denoted the change of d-score as Δ_all_d-score (Figure 4C). Interestingly, alterations of methylation states on different CpG sites contributed differently to the change of d-scores (Figure 4D), which couldn’t be fully explained by the alteration of methylation ratios. Besides, when we altered reads from a whole methylated to a whole unmethylated state, though with the same alteration of methylation ratios, changes of d-scores varied across the region (Figure 4E), which was not entirely determined by the amount of CpG sites on reads. All the results showed that DISMIR assigned different importance to different CpG sites according to their surrounding DNA sequences. We further added all Δ_single_d-scores of a read together as Δ_sum_d-score, and found that almost all Δ_sum_d-scores were less than the corresponding Δ_all_d-scores (Figure 4E). The results suggested that DISMIR may focus more on the global methylation alteration rather than just gather the impact of each single CpG site’s methylation alteration together.

### Motif-related kernels of DISMIR could resist to methylation state alterations of single CpG sites

As shown in Figure 4F, the d-score change with the alteration of all CpG sites on a read was much higher than the sum of d-score changes with alterations of each single CpG site. This result hinted that DISMIR might pay more attention to global methylation state alterations which are familiar in cancer tissues. By contrast, alterations of single CpG sites are usually confounded by technical noise during WGBS, thus should be considered with smaller weights when discriminating the origin of reads. To further investigate whether DISMIR could resist to methylation state alterations of single CpG sites, we considered all possible reads with more than three CpG sites in all switching regions and calculated their Δ_all_d-scores and Δ_sum_d-scores. To avoid the confounding of the counts of CpG sites, we grouped the reads by the CpG count and analyzed each group respectively. Interestingly, DISMIR paid more attention to global alteration of CpG states beyond the additive model of single alterations (the left panel of Figure 5A showed results of all reads with three CpG sites; reads with more CpG sites showed similar patterns). The results suggested DISMIR worked as a filter against the influence of the methylation alterations on single CpG sites which have low signal-to-noise ratios in comparison with global methylation alterations.

**Figure 5.**
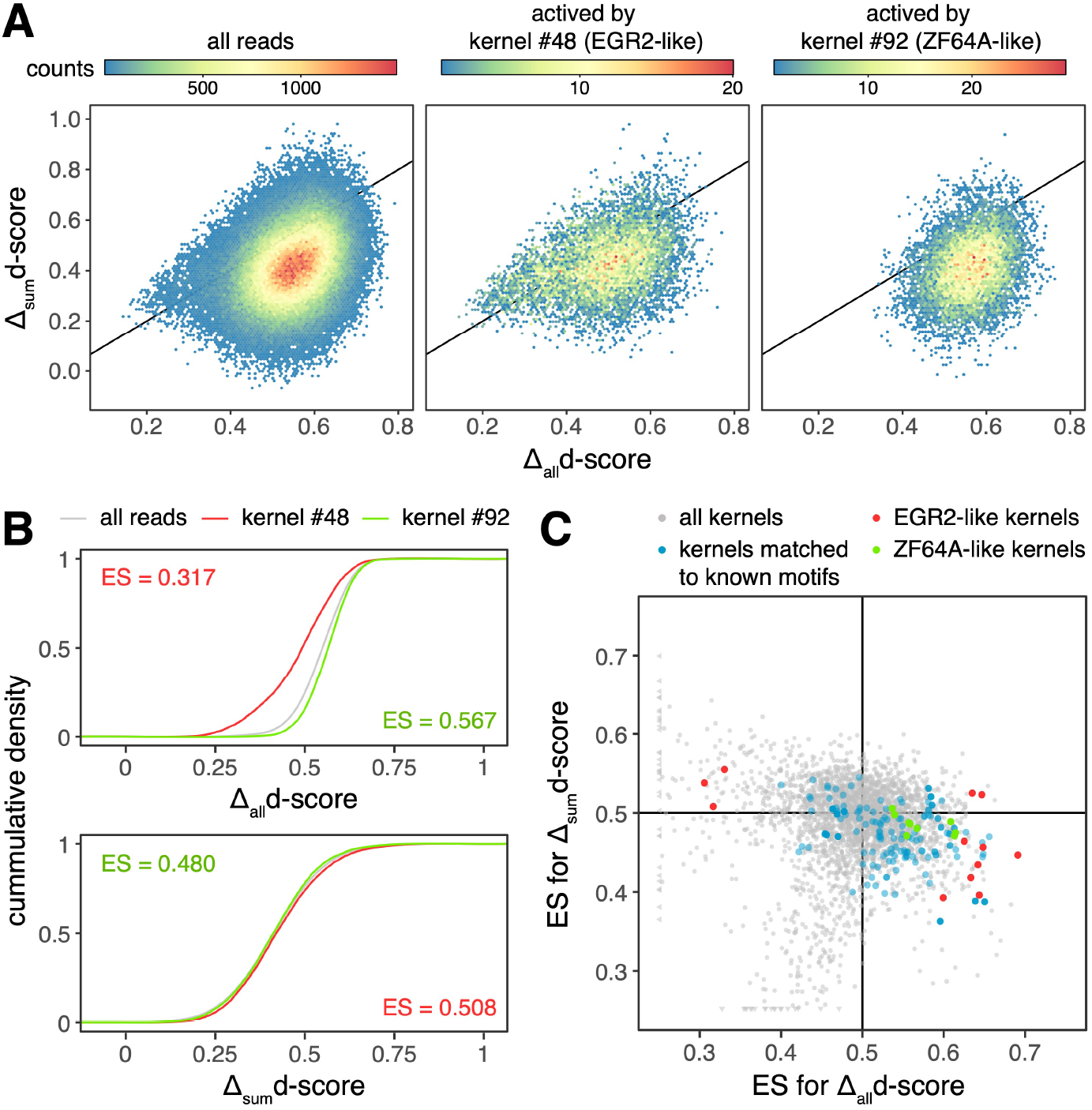
The ability of DISMIR and its kernels on resisting to methylation state alterations of single CpG sites. (A) The distribution of Δ_all_d-scores and Δ_sum_d-scores of all reads (left) and certain kernel-activated reads (middle and right). The black lines are diagonal lines where Δ_all_d-scores are equal to Δ_sum_d-scores. (B) The cumulative distribution function of Δ_all_d-scores and Δ_sum_d-scores of reads shown in (A). Effect sizes (ES) of the Mann–Whitney *U* test are shown. (C) The distribution of ES of the Mann–Whitney *U* test for Δ_all_d-scores and Δ_sum_d-scores of reads activated by certain kernels in comparison with them of all reads, which was performed the same as shown in (B). For (A) and (B), results of reads with three CpG sites were shown. For (C), results of reads with three, four and five CpG sites were shown with performing model training for ten times. Black lines show where ES equals to 0.5 and divide the plane into four quadrants.

We then analyzed the relationship between d-score changes and DISMIR kernels. We calculated the activation values of each kernel on every possible read with all CpG sites set as methylated, and reads with the top 0.1% highest activation values were regarded as reads that could highly activate this kernel. Interestingly, the distribution of d-score changes of kernel-activated reads tended to be different from the distribution of all reads (Figure 5A & B). We further compared the distribution of Δ_all_d-scores and Δ_sum_d-scores of these kernel-activated reads and all reads with the Mann–Whitney *U* test. As sample sizes of both distributions were huge, the *p*-value of the test was overpowered for subtle difference. Therefore, we adopted the AUC statistic between two distributions, which could be directly derived from the Mann–Whitney *U* test (Mason and Graham, 2002), as the effect size (ES) of the test to quantify the difference between two distributions. As shown in Figure 5C, some kernels tended to filter out the influence of single site demethylation but could focus on whole demethylation (located in the fourth quadrant of Figure 5C). We further investigated kernels matched to known motifs (blue points in Figure 5C) and found that these kernels are more likely to be located in the fourth quadrant (odds ratio = 4.158, Fisher’s exact test *p*-value = 4.59×10^−23^). As the randomness of the training of the deep learning model, kernels may differ across different training, but functional kernels which are more likely to be assigned to known motifs emerged repeatedly in different trainings. These functional kernels, as shown in Figure 5C, could resist to the demethylation of single CpG sites, ensuring the high robustness of DISMIR.

## DISCUSSION

In this study, we developed a deep learning-based approach called DISMIR to predict whether reads in plasma cfDNA WGBS data are derived from tumor, and further adopted the predicted fraction of tumor-derived reads to diagnose cancer. DISMIR achieved outperformed results in HCC detection, especially at low sequencing depths, which makes it possible to be a low-cost cancer-detection method. The predicted fractions of tumor-derived reads are also stable at different sequencing depths, so that we can assign a unified threshold to samples with various sequencing depths for cancer diagnosis. These advantages make DISMIR more likely to be applied in clinical practice.

The outperformance of DISMIR was mainly contributed by the novel design of the deep learning model. We built a deep network to combine the DNA sequence and methylation information together for each read. Therefore, DISMIR could grasp sequence motifs related to cancer and extract the joint patterns of DNA sequence and methylation across different regions from the whole genome to ensure the source prediction of individual reads more accurate. Information derived from different regions makes the model more robust and thus guarantees the precision of prediction even at extremely low sequencing depths.

Different to the definitions of DMRs in other approaches, here we introduced a novel method to identify DMRs called switching regions to enrich reads with more distinguishable methylation patterns. As previous studies suggested, methylation patterns at the resolution of read level could make the model more sensitive (Lee *et al.*, 2019). DNA fragments from switching regions contain more specific features, and could thus enhance the precision of signal detection resisting to noise at low sequencing depths. Thus, defining DMRs by switching regions is more suitable for models employing individual reads as inputs. Furthermore, kernels of DISMIR that were related to known motifs paid more attention to global alteration of methylation states but less attention to methylation state alterations of single CpG sites, which made DISMIR able to resist to technical noise of WGBS, and thus enhanced the robustness of DISMIR.

We found that several motifs that kernels of the deep learning model focused on were related to cancer, which showed the powerful capacity of feature extraction as well as good interpretability of the deep learning model. Furthermore, some kernels that didn’t match with known cancer motifs may contain novel information related to cancer, especially in the process of epigenomic regulation. The deep learning method, which integrates DNA sequence and methylation information together, also provides a data-driven method for us to unveil the interaction between genomes and epigenomes (Eraslan *et al.*, 2019; Zou *et al.*, 2019). In addition, the method can also be applied in other kinds of data. Deep learning model can integrate different forms or diverse information levels of data together to extract useful joint patterns to find out new rules in certain biological processes.

This study can be further improved in several ways. Firstly, more cfDNA samples could be involved in the testing cohort to further evaluate the precision of the method. Besides, as the tumor samples used for model training may not be just composed of cancer cells (Aran *et al.*, 2015), we can develop correction methods based on the tumor purity before model training to get more accurate prediction of tumor-derived cfDNA. Furthermore, though this study employed HCC to evaluate the performance, the method could be used and should be validated on more kinds of cancers. In addition, the method could be extended from two-source prediction to multiple-source prediction, and thus realize pan-cancer detection.

## SUPPLEMENTARY DATA

Supplemental Table 1 Supplemental codes of DISMIR

## ACKNOWLEDGEMENT

The authors greatly acknowledge Prof. Yuk Ming Dennis Lo and Prof. Peiyong Jiang for sharing cfDNA WGBS data (Chan *et al.*, 2013).

## REFERENCES

Abbosh,C. et al. (2017) Phylogenetic ctDNA analysis depicts early-stage lung cancer evolution. Nature,545, 446–451.

Adalsteinsson,V.A. et al. (2017) Scalable whole-exome sequencing of cell-free DNA reveals high concordance with metastatic tumors. Nature Communications,8.

Alvarez,H. et al. (2011) Widespread hypomethylation occurs early and synergizes with gene amplification during esophageal carcinogenesis. PLoS Genet.,7, e1001356.

Aran,D. et al. (2015) Systematic pan-cancer analysis of tumour purity. Nat Commun,6, 8971.

Baylin,S.B. et al. (2001) Aberrant patterns of DNA methylation, chromatin formation and gene expression in cancer. Hum. Mol. Genet.,10, 687–692.

Bettegowda,C. et al. (2014) Detection of circulating tumor DNA in early- and late-stage human malignancies. Sci Transl Med,6, 224ra24.

Bitzer,M. et al. (2016) Resminostat plus sorafenib as second-line therapy of advanced hepatocellular carcinoma – The SHELTER study. Journal of Hepatology,65, 280–288.

Burrell,R.A. et al. (2013) The causes and consequences of genetic heterogeneity in cancer evolution. Nature,501, 338–345.

Cedar,H. and Bergman,Y. (2012) Programming of DNA Methylation Patterns. Annu. Rev. Biochem.,81, 97–117.

Chan,K.C.A. et al. (2013) Noninvasive detection of cancer-associated genome-wide hypomethylation and copy number aberrations by plasma DNA bisulfite sequencing. Proceedings of the National Academy of Sciences,110, 18761–18768.

Chicard,M. et al. (2016) Genomic Copy Number Profiling Using Circulating Free Tumor DNA Highlights Heterogeneity in Neuroblastoma. Clin Cancer Res,22, 5564–5573.

Cristiano,S. et al. (2019) Genome-wide cell-free DNA fragmentation in patients with cancer. Nature,570, 385–389.

Crowley,E. et al. (2013) Liquid biopsy: monitoring cancer-genetics in the blood. Nature Reviews Clinical Oncology,10, 472–484.

Eraslan,G. et al. (2019) Deep learning: new computational modelling techniques for genomics. Nat Rev Genet,20, 389–403.

Feinberg,A.P. et al. (2006) The epigenetic progenitor origin of human cancer. Nat. Rev. Genet.,7, 21–33.

Feng,H. et al. (2019) Disease prediction by cell-free DNA methylation. Briefings in Bioinformatics,20, 585–597.

Guo,W. et al. (2013) BS-Seeker2: a versatile aligning pipeline for bisulfite sequencing data. BMC Genomics,14, 774.

Gupta,S. et al. (2007) Quantifying similarity between motifs. Genome Biol.,8, R24.

Hebestreit,K. et al. (2013) Detection of significantly differentially methylated regions in targeted bisulfite sequencing data. Bioinformatics,29, 1647–1653.

Heitzer,E. et al. (2018) Current and future perspectives of liquid biopsies in genomics-driven oncology. Nature Reviews Genetics.

Jühling,F. et al. (2016) metilene: fast and sensitive calling of differentially methylated regions from bisulfite sequencing data. Genome Res.,26, 256–262.

Lee,D. et al. (2019) PRISM: methylation pattern-based, reference-free inference of subclonal makeup. Bioinformatics,35, i520–i529.

Li,S. et al. (2013) An optimized algorithm for detecting and annotating regional differential methylation. BMC Bioinformatics,14 Suppl 5, S10.

Li,W. et al. (2018) CancerDetector: ultrasensitive and non-invasive cancer detection at the resolution of individual reads using cell-free DNA methylation sequencing data. Nucleic Acids Research.

Lienert,F. et al. (2011) Identification of genetic elements that autonomously determine DNA methylation states. Nat Genet,43, 1091–1097.

Liggett,T. et al. (2010) Differential methylation of cell-free circulating DNA among patients with pancreatic cancer versus chronic pancreatitis. Cancer,116, 1674–1680.

Mason,S.J. and Graham,N.E. (2002) Areas beneath the relative operating characteristics (ROC) and relative operating levels (ROL) curves: Statistical significance and interpretation. Q. J. R. Meteorol. Soc.,128, 2145–2166.

Newman,A.M. et al. (2014) An ultrasensitive method for quantitating circulating tumor DNA with broad patient coverage. Nat Med,20, 548–554.

Quang,D. and Xie,X. (2016) DanQ: a hybrid convolutional and recurrent deep neural network for quantifying the function of DNA sequences. Nucleic Acids Res.,44, e107.

Schwarzenbach,H. et al. (2011) Cell-free nucleic acids as biomarkers in cancer patients. Nature Reviews Cancer,11, 426–437.

Shao,C. et al. (2009) Hemimethylation footprints of DNA demethylation in cancer. Epigenetics,4, 165–175.

Snyder,M.W. et al. (2016) Cell-free DNA Comprises an In Vivo Nucleosome Footprint that Informs Its Tissues-Of-Origin. Cell,164, 57–68.

Wan,J.C.M. et al. (2017) Liquid biopsies come of age: towards implementation of circulating tumour DNA. Nature Reviews Cancer,17, 223–238.

Wang,J. et al. (2020) NFAT2 overexpression suppresses the malignancy of hepatocellular carcinoma through inducing Egr2 expression. BMC Cancer,20, 966.

Warton,K. and Samimi,G. (2015) Methylation of cell-free circulating DNA in the diagnosis of cancer. Front. Mol. Biosci.,2.

Weiss,G.J. et al. (2017) Tumor Cell–Free DNA Copy Number Instability Predicts Therapeutic Response to Immunotherapy. Clin Cancer Res,23, 5074–5081.

Wu,H. et al. (2015) Detection of differentially methylated regions from whole-genome bisulfite sequencing data without replicates. Nucleic Acids Res.,43, e141.

Zeng,T. et al. (2017) LncRNA-AF113014 promotes the expression of Egr2 by interaction with miR-20a to inhibit proliferation of hepatocellular carcinoma cells. PLoS ONE,12, e0177843.

Zou,J. et al. (2019) A primer on deep learning in genomics. Nat Genet,51, 12–18.

