## Supplemental Table 1 for "DISMIR: a deep learning-based cancer-detection method by integrating DNA sequence and methylation information of individual cell-free DNA reads"

**SUPPLEMENTAL MATERIALS**

Supplemental Table 1. Significant motifs (*p*-value < 0.05) matched with the kernel PFMs. *E*-values assigned by TOMTOM from ten times of training were merged with Fisher's combined probability test and the *p*-values were shown. References of studies that provided the evidence between the motif and HCC were supplemented.

| Motif | *p*-value | Ref. |
|---|---|---|
| ZBT17 | $2.446\times10^{-7}$ | |
| SP2 | $4.516\times10^{-6}$ | (1) |
| SP3 | $2.276\times10^{-5}$ | (2) |
| EGR2 | $7.754\times10^{-5}$ | (3, 4) |
| ZFX | $1.549\times10^{-4}$ | (5) |
| MXI1 | $1.699\times10^{-4}$ | (6) |
| ZF64A | $3.215\times10^{-4}$ | (7) |
| VEZF1 | $7.591\times10^{-4}$ | |
| ZN341 | $9.327\times10^{-4}$ | |
| GATA1 | $1.242\times10^{-3}$ | (8) |
| MAZ | $1.670\times10^{-3}$ | (9) |
| ZN263 | $2.541\times10^{-3}$ | |
| PATZ1 | $2.758\times10^{-3}$ | (10) |
| RARA | $3.501\times10^{-3}$ | (11) |

| Motif | *p*-value | Ref. |
|---|---|---|
| WT1 | $4.346\times10^{-3}$ | (12) |
| KLF16 | $5.402\times10^{-3}$ | |
| RARG | $5.627\times10^{-3}$ | (13) |
| SP1 | $6.093\times10^{-3}$ | (14) |
| PLAG1 | $1.002\times10^{-2}$ | (15) |
| EGR1 | $1.020\times10^{-2}$ | (16) |
| TAL1 | $1.298\times10^{-2}$ | (17) |
| E2F4 | $1.476\times10^{-2}$ | (18) |
| ZN467 | $1.794\times10^{-2}$ | |
| TBX15 | $2.255\times10^{-2}$ | (19) |
| SMAD3 | $3.128\times10^{-2}$ | (20) |
| KLF6 | $3.909\times10^{-2}$ | (21) |
| ZN770 | $4.691\times10^{-2}$ | |
| HINFP | $4.705\times10^{-2}$ | (22) |


1. Zhu,Y., Cui,J., Liu,J., Hua,W., Wei,W. and Sun,G. (2020) Sp2 promotes invasion and metastasis of hepatocellular carcinoma by targeting TRIB3 protein. *Cancer Med*, **9**, 3592–3603.

2. Huang,Z., Huang,L., Shen,S., Li,J., Lu,H., Mo,W., Dang,Y., Luo,D., Chen,G. and Feng,Z. (2015) Sp1 cooperates with Sp3 to upregulate MALAT1 expression in human hepatocellular carcinoma. *Oncology Reports*, **34**, 2403–2412.

3. Zeng,T., Wang,D., Chen,J., Tian,Y., Cai,X., Peng,H., Zhu,L., Huang,A. and Tang,H. (2017) LncRNA-AF113014 promotes the expression of Egr2 by interaction with miR-20a to inhibit proliferation of hepatocellular carcinoma cells. *PLoS ONE*, **12**, e0177843.

4. Wang,J., Zhang,Y., Liu,L., Cui,Z., Shi,R., Hou,J., Liu,Z., Yang,L., Wang,L. and Li,Y. (2020) NFAT2 overexpression suppresses the malignancy of hepatocellular carcinoma through inducing Egr2 expression. *BMC Cancer*, **20**, 966.

5. Ding,W., Tan,H., Li,X., Zhang,Y., Fang,F., Tian,Y., Li,J. and Pan,X. (2018) MicroRNA-493 suppresses cell proliferation and invasion by targeting ZFX in human hepatocellular carcinoma. *CBM*, **22**, 427–

434.

6. Sharma,B.K., Kolhe,R., Black,S.M., Keller,J.R., Mivechi,N.F. and Satyanarayana,A. (2016) Inhibitor of differentiation 1 transcription factor promotes metabolic reprogramming in hepatocellular carcinoma cells. *FASEB j.*, **30**, 262–275.

7. Bitzer,M., Horger,M., Giannini,E.G., Ganten,T.M., Wörns,M.A., Siveke,J.T., Dollinger,M.M., Gerken,G., Scheulen,M.E., Wege,H., *et al.* (2016) Resminostat plus sorafenib as second-line therapy of advanced hepatocellular carcinoma – The SHELTER study. *Journal of Hepatology*, **65**, 280–288.

8. Andrieux,L.O., Fautrel,A., Bessard,A., Guillouzo,A., Baffet,G. and Langouët,S. (2007) GATA-1 Is Essential in EGF-Mediated Induction of Nucleotide Excision Repair Activity and ERCC1 Expression through ERK2 in Human Hepatoma Cells. *Cancer Res*, **67**, 2114–2123.

9. Luo,W., Zhu,X., Liu,W., Ren,Y., Bei,C., Qin,L., Miao,X., Tang,F., Tang,G. and Tan,S. (2016) MYC associated zinc finger protein promotes the invasion and metastasis of hepatocellular carcinoma by inducing epithelial mesenchymal transition. *Oncotarget*, **7**, 86420–86432.

10. Valentino,T., Palmieri,D., Vitiello,M., Pierantoni,G.M., Fusco,A. and Fedele,M. (2013) PATZ1 interacts with p53 and regulates expression of p53-target genes enhancing apoptosis or cell survival based on the cellular context. *Cell Death Dis*, **4**, e963–e963.

11. Sano,K., Takayama,T., Murakami,K., Saiki,I. and Makuuchi,M. (2003) Overexpression of retinoic acid receptor alpha in hepatocellular carcinoma. *Clin Cancer Res*, **9**, 3679–3683.

12. Mžik,M., Chmelařová,M., John,S., Laco,J., Slabý,O., Kiss,I., Bohovicová,L., Palička,V. and Nekvindová,J. (2016) Aberrant methylation of tumour suppressor genes WT1, GATA5 and PAX5 in hepatocellular carcinoma. *Clinical Chemistry and Laboratory Medicine (CCLM)*, **54**.

13. Gan,W.-J., Wang,J.-R., Zhu,X.-L., He,X.-S., Guo,P.-D., Zhang,S., Li,X.-M., Li,J.-M. and Wu,H. (2016) RARγ-induced E-cadherin downregulation promotes hepatocellular carcinoma invasion and metastasis. *J Exp Clin Cancer Res*, **35**, 164.

14. Zhang,X., Zhuang,H., Han,F., Shao,X., Liu,Y., Ma,X., Wang,Z., Qiang,Z. and Li,Y. (2018) Sp1-regulated transcription of RasGRP1 promotes hepatocellular carcinoma (HCC) proliferation. *Liver Int*, **38**, 2006–2017.

15. Cao,Y., Tao,Q., Kao,X. and Zhu,X. (2020) Hsa-circRNA-103809 Promotes Hepatocellular Carcinoma Development via MicroRNA-1270/PLAG1 Like Zinc Finger 2 Axis. *Dig Dis Sci*, 10.1007/s10620-020-06416-x.

16. Zhang,Q., Song,G., Yao,L., Liu,Y., Liu,M., Li,S. and Tang,H. (2018) miR-3928v is induced by HBx via NF-κB/EGR1 and contributes to hepatocellular carcinoma malignancy by down-regulating VDAC3. *J Exp Clin Cancer Res*, **37**, 14.

17. Zou,R.-C., Xiao,S.-F., Shi,Z.-T., Ke,Y., Tang,H.-R., Wu,T.-G., Guo,Z.-T., Ni,F., Li,W.-X. and Wang,L. (2019) Identification of metabolism-associated pathways and genes involved in male and female liver cancer patients. *J Theor Biol*, **480**, 218–228.

18. Yang,W.-X., Pan,Y.-Y. and You,C.-G. (2019) CDK1, CCNB1, CDC20, BUB1, MAD2L1, MCM3, BUB1B, MCM2, and RFC4 May Be Potential Therapeutic Targets for Hepatocellular Carcinoma Using Integrated Bioinformatic Analysis. *BioMed Research International*, **2019**, 1–16.

19. Zheng,Y., Huang,Q., Ding,Z., Liu,T., Xue,C., Sang,X. and Gu,J. (2016) Genome-wide DNA methylation analysis identifies candidate epigenetic markers and drivers of hepatocellular carcinoma. *Brief Bioinform*, 10.1093/bib/bbw094.

20. Fu,Q., Zhang,Q., Lou,Y., Yang,J., Nie,G., Chen,Q., Chen,Y., Zhang,J., Wang,J., Wei,T., *et al.* (2018) Primary tumor-derived exosomes facilitate metastasis by regulating adhesion of circulating tumor cells via SMAD3 in liver cancer. *Oncogene*, **37**, 6105–6118.

21. Kremer-Tal,S., Reeves,H.L., Narla,G., Thung,S.N., Schwartz,M., Difeo,A., Katz,A., Bruix,J., Bioulac-Sage,P., Martignetti,J.A., *et al.* (2004) Frequent inactivation of the tumor suppressor Kruppel-like factor 6 (KLF6) in hepatocellular carcinoma. *Hepatology*, **40**, 1047–1052.

22. Zhu,Y.-Z., Zhu,R., Fan,J., Pan,Q., Li,H., Chen,Q. and Zhu,H.-G. (2010) Hepatitis B virus X protein induces hypermethylation of p16 $^{INK4A}$ promoter via DNA methyltransferases in the early stage of HBV-associated hepatocarcinogenesis. *Journal of Viral Hepatitis*, **17**, 98–107.
